# A sigmoid curve analysis method for pharmacological experimental results

**DOI:** 10.1101/2022.12.18.520530

**Authors:** Qingxia Niu, Chengyan Zhao

**Affiliations:** Department of Pathophysiology, Shantou University Medical College, Shantou, 515041, Guangdong, China

**Keywords:** Sigmoid curve, Sigmoid equation, Sigmoid curve analysis method, Sigmoid curve linear fitting, ED_50_, pharmacology, mass action law, hyperbolic equation

## Abstract

Sigmoid curve (S-curve) is a basic exhibition form of dose-effect relationship in drug reaction. To analyze S-curve is an important method to well-understand drug reaction performance (DRP). The present study introduced an S-curve analysis method for pharmacological experiment results (PERs), the core of which was to solve the problem of the linear fitting of S-curve equation (S-Eq). The linear fitting Eqs of S-Eq were established with 100% fitness. Meanwhile, mathematical and pharmacological meaning of S-curve constants, ED_50_ and maximum effect (y_max_) were clarified. The same group of experimental data was analyzed by the present method and four traditional analysis methods. The result indicates that the experimental parameters and their values displaying DRP got by different methods are different. The S-curve analysis method is closer to real drug reaction law.

## 1. Introduction

Many biological phenomena frequently exhibit S-curve, such as time-effect relationship of cell/microbial proliferation^1,2^ and dose-effect relationship shown in antigen-antibody reaction^3^, receptor-ligand reaction^4^, cytotoxic reaction^5^, enzyme-substrate reaction^6,7^, and drug-target reaction^8,9^. S-curve is the basic exhibition form of dose-effect relationship in PERs. Not all PERs can exhibit typical S-curve because of experimental methods and conditions. Most of PERs only exhibit a part of S-curve. To analyze S-curve is an important method to study and well-understand biological reaction law. It is essential that to analyze S-curve needs to convert S-Eq into linear Eq and to clarify pharmacological meanings of S-Eq constants.

S-Eq is S-curve mathematical expression. Up to now, to analyze PERs have employed the traditional mathematical Eqs instead of classical S-Eq, the major reason of which is perhaps that a way to solve S-Eq with three constants (a, b, and k) has not been yet found^10-13^. The traditional analysis method takes the partial exhibition form of PERs as a whole, such as exponential (exp), logarithmic (lg), double logarithmic (dlg) or hyperbolic curve^14^. The approximate Eqs to S-Eq are used to analyze PERs, which could not reflect accurate drug reaction law. This study established the linear fitting method for basic S-Eq and clarified the mathematical and pharmacological meaning of S-Eq constants. The combination of the two makes the S-Eq hard problem be solved. The clarification and application of the present analysis method is of great value for both pharmacological and biological study.

## 2. The relationship between PERs and S-curve

### 2.1. S-curve is the basic exhibition form of dose-effect relationships produced by drug reactions

Pharmacology includes pharmacokinetics and pharmacodynamics. Drug reaction belongs to chemical reaction. Chemical reaction follows mass action law (MAL). Many literatures define MAL as: Chemical reaction velocity (*ν*) is directly proportional to the effective mass of reactants. Effective mass refers to molecule number per unit volume, *i.e*., gas density or liquid density (concentration), but no absolute mass. This definition is actually in dispute.

In 1867, MAL was pinpointed in the book《Research on chemical affinity》by a mathematician Guldberg CM and a chemist Waage P after they carefully studied the interaction principle of substances in the process of chemical reaction. They believe that the most natural way to study chemical reactions is to determine force balance. The force to produce new compounds can interact with other forces in chemical reaction. The reaction is reversible, incomplete, and finally reaches equilibrium. This force is called as affinity. For example, NO_2_+CO reaction generates CO_2_+NO, but CO_2_ and NO reaction also products NO_2_ and CO at the same time.

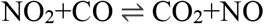

At reaction equilibrium, the affinity is directly proportional to the effective mass product of reactants. [CO_2_], [NO], [NO_2_] and [CO] represent CO_2_, NO, NO_2_ and CO concentrations, respectively. *k* stands for affinity coefficient. MAL can be shown as follows.

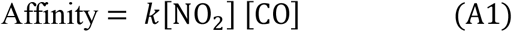

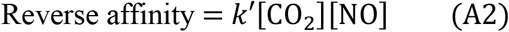

The two affinities are equal when reaction balance is reached, then *k*[NO_2_] [CO]= *k*′[CO_2_][NO]. If k stands for total affinity coefficient, and k = *k/k*′ = [CO_2_][NO]/[NO_2_] [CO], then

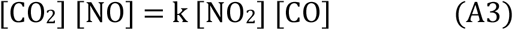

It is probable because the meaning of affinity and its coefficient were not clearly explained. In 1877, van’t Hoff, a Dutch chemist, restated MAL as *ν* is directly proportional to the effective mass product of reactants. In the definition, the affinity with unclear meaning is substituted by *ν*.

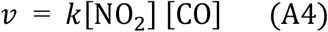

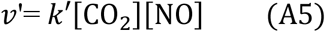

van’t Hoff believed that total v = *ν*-*ν*’= *k*[NO_2_] [CO] − *k*′[CO_2_][NO]. When the reaction reaches equilibrium, v=0, then, *k*[NO_2_] [CO] = *k*′[CO_2_][NO].

Let k=*k/k*′= [CO_2_][NO]/[NO_2_][CO], then, [CO_2_][NO]=k[NO_2_][CO].

The result is the same as Eq. (A3). Concentration units are usually mmol·mL^-1^, mol/L^-1^, etc. Time units are s, min, h, etc. Velocity units are available in mmol·mL^-1^·s^-1^, mol/L^-1^·min^-1^, etc.

The velocity is clear in meaning, that is, the effective mass or concentration of reaction products per unit time. Velocity includes immediate *ν* and average velocity 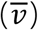. Generally, 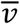 is measured, that is, product concentrations are measured in a fixed time. As above reaction, the product can be CO_2_ and NO. The two product concentrations are the same. The CO_2_ and NO amount are related to measurement time. Long time is, more product amount is, and short time is, less product amount is, when the reaction does not reach plateau (*ν*≠0). So, CO_2_ and NO amount are dependent on the selected time period. 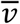 is invariableness. The concentrations of reactant and product are stable when the reaction reaches plateau. The production of CO_2_ and NO no longer increases with time at 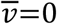. That is, the product amount is independent of *ν*, which is inconsistent with the definition of MAL by van’t Hoff. Obviously, it is inappropriate that he defined MAL by replacing affinity with *ν*. In the definition of Guldberg CM and Waage P, affinity is reflected by its coefficient k that is clear in meaning. The k value can be determined by experiments. The correct definition for MAL should be indicated by Eq. (A3): The effective mass product of products is directly proportional to the effective mass product of reactants, which is unrelated to *ν*.

If Eq. (A3) is transformed, then an analytical form of the experimental results can be got.

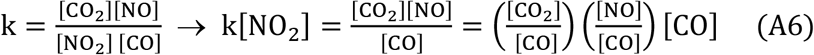

Here, [CO_2_]/[CO] and [NO]/[CO] are the growth ratio (GR) of the two products (CO_2_ and NO). Because [CO_2_] = [NO], or [CO_2_]/[CO] = [NO]/[CO], Eq. (A6) can be transformed into Eq. (A7).

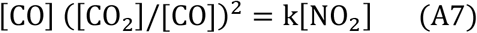

Generally, one plays a dominant role, and the other plays passive role when two substances react. Enzyme-substrate reaction is the typical case^14,15^. Enzyme plays a dominant role. Substrate, the subject by action, catalyzed by enzyme, forms products. Pharmacological experiment usually takes drug concentrations (doses) as independent variables (*x*) which react with a certain amount of substrate. Product growth ratio (GR) or substrate depletion ratio (DR) is used as an effect indicator (dependent variables, function, or *y*). Thus, DRP can be understood by analyzing the relationship between *x* and *y*.

Eq. (A7) is the dose-effect relationship of the reaction system consisting of two reactants and two products. If the reaction system consists of two reactants and one product, such as receptor-ligand reaction, then the product only possesses the receptor-ligand conjugate^16^. In this case, Eq. (A7) can be transformed into [CO_2_]/[CO] = k[NO_2_]. If the reaction system is two reactants and three products, or multiple products, then Eq. (A7) can be transformed into Eq. (A8).

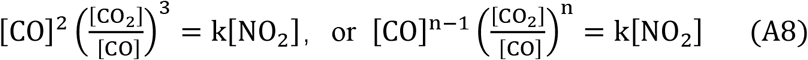

A simple and intuitive linear Eq. (A9) can be obtained when *y* = [CO]^n−1^([CO_2_] /[CO])^n^ and *x* = [NO_2_] are put into Eq. (A8).

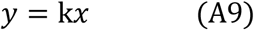

Eqs. (A8 and A9) are another expression of MAL. That is, the product of GR n-power of reaction products and the (n-1)-power of substrate concentrations is proportional to reactant concentrations.

MAL is derived from simple experimental results completed by one step reaction in low density gas and “dilute solution” matter. It is only suitable for the simple reaction completed by one step in “low-density” gas and “dilute solution” matter, but unsuitable for any other chemical reaction in biological experiments. The quantification bounds of “low density” or “dilute solution” depend on reactant properties and need to be determined experimentally. Pharmacological study is performed in reaction system at set concentrations. The reaction follows MAL if the effective mass of reactants is within the limited “low-density” and “dilute solution” range. If the experiment condition is below the lower limit or above the upper limit for the limited range, the reaction certainly disobeys MAL. What law does the reaction follow?

Based on the derivation principle of MAL, affinity plays a main role in chemical reaction. affinity size is inversely proportional to the distance square between reactants. It is well-known that the mutual contact between reactants is a necessary condition, suggesting that distance is the main factor affecting the reaction^17^. Below the lower limit of the “low-density” or “dilute solution” range, the distance between reactants gradually approached the limited range from far to near, the contact probability of reactants increased from very low levels. The product increased with reactant density rise, affinities increased exponentially, and the dose-effect relationship showed an exp curve (part I). Under low-density” and “dilute solution” conditions, reactants can fully contact each other, so that the reaction followed MAL. In this case, the dose-effect relationship exhibited a straight line (part II). Above the upper limit of the “low-density” or “dilute solution” range, the product lowly increased with the reactant density rise, which prevents reactants from the contact with each other, because the contact probability decreased. The dose-effect relationship showed a curve elevating slowly (part III). When the dose-effect curve (DEC) for the entire reaction was connected the plot on the two-dimensional coordinate was an S-curve located in quadrants I and II (Fig. 1). This is consistent with the DEC obtained from pharmacological experiments. S-curve not only showed the chemical reaction following MAL, but also showed the chemical reaction disobeying MAL. Accordingly, the PERs expressed by the dose-effect relationship should be regarded as S-curves for analysis. The experimental conditions set are not all necessarily appropriate. Thus, not all PERs exhibit complete S-curves. Some of PERs are only a part of S-curve, such as near-exp, near-lg curve, which should also be regarded as S-curve to be analyzed.

**Figure 1.**
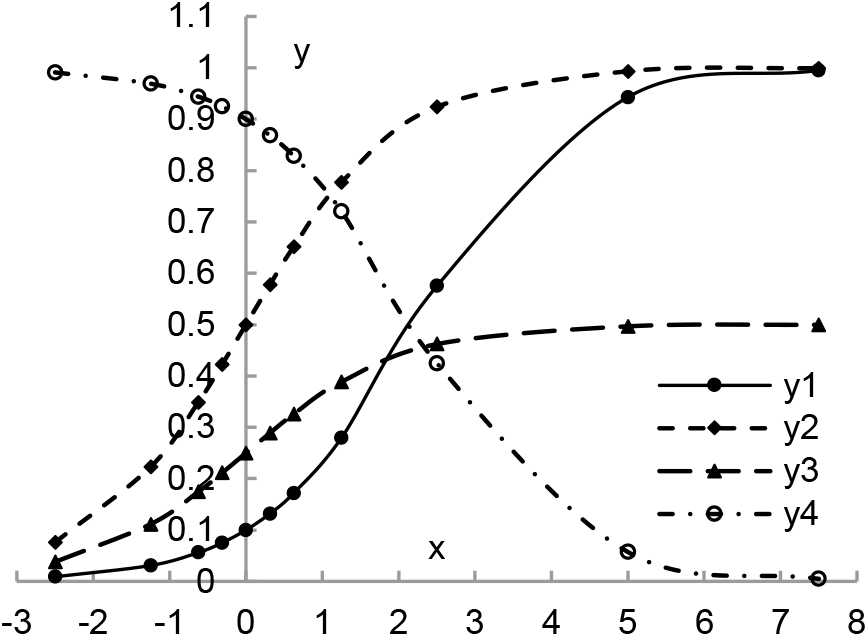
Standard S-curves derived from S-Eqs. (S1 and S2). When different constant values were put into S-Eqs. (S1 and S2) y values can be calculated. Four S-curves are plotted. y_1_, y_2_ and y_3_ are rising curves. y_4_ (an anti-function of y_1_ at r=1) is decreasing curve.

Eq. (A9) is a linear Eq, not an S-curve, expressing the dose-effect relationship with a straight line. However, actual drug reaction process exists as S-curve, which requires the Eq expressing S-curve. There are several S-Eqs in mathematical field. A basic S-Eq that is Eq. (S1) and its reverse function, Eq. (S2), was employed in the present study. The function of Eq. (A9) is contained in S-Eq. (S1).

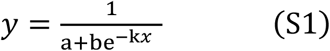

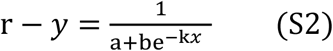

Here, *x, y*, and k are doses (independent variables), effect or effect indicators (dependent variables), and affinity constant, respectively. *x* and *y* in Eq. (S1) have the same meaning as *x* and *y* in Eq. (A9). Constants (a, b, and k) determine S-curve shape. S-Eqs. (S1 and S2) have the following characteristics. ①When *x*→∝, *y* in Eq. (S1) has maximal value (y_max_). y_max_ = 1/a. When *x*=0, y_0_ = 1/(a + b). y_0_ is S-curve intercept. ②When *x*→∝, *y* in Eq. (S2) has minimal value, y=0. When y=0 enters Eq. (S2) r = 1/a.

### 2.2. Pharmacological meanings of S-Eq constants

An experimental system for drug reaction is usually established in pharmacological experiments. Different drug dose can produce different effects. Different dose (*x*)-effect (*y*) relationship forms different S-curve shape that is determined by S-Eq constant size. Fig. 2 shows the relationship between S-curve shape and constant values, which reveal the pharmacological meanings of S-Eq constants.

**Figure 2.**
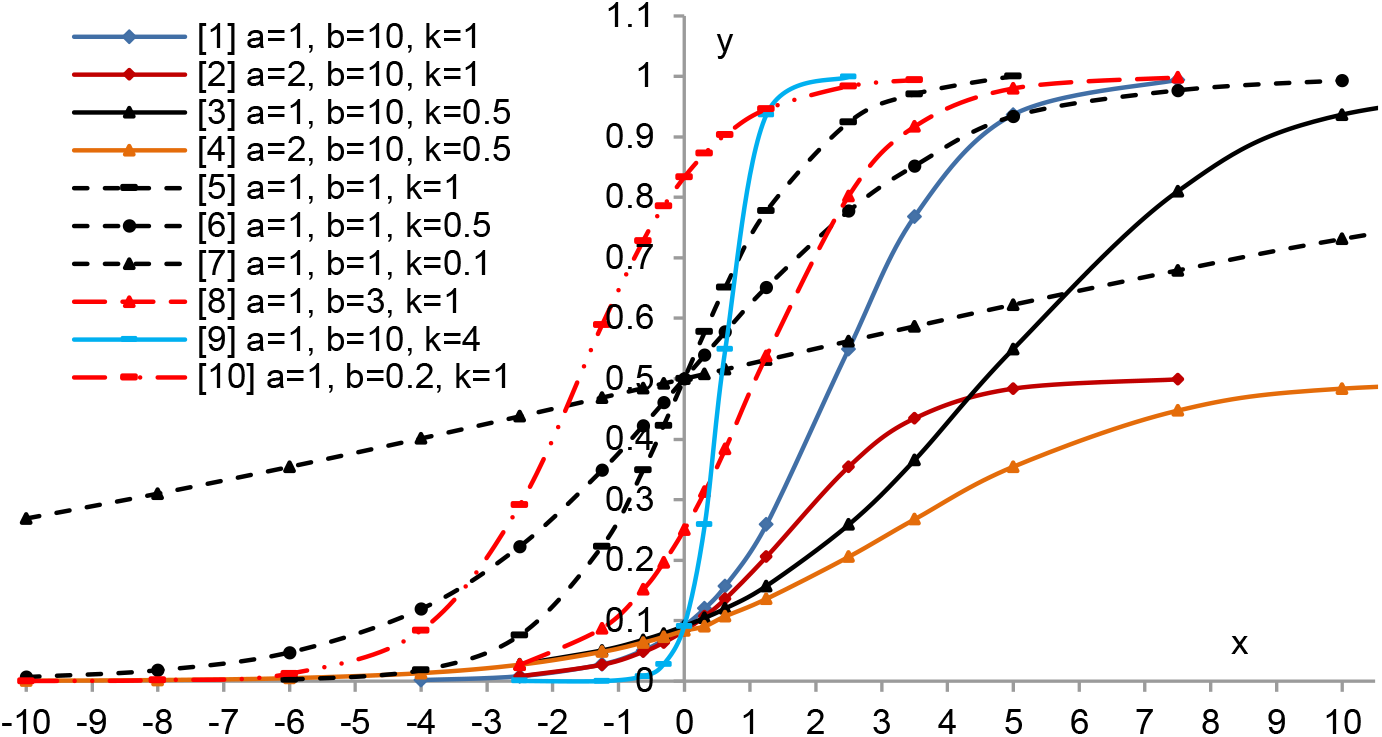
The relationship between S-Eq constants and S-curve shape plotted by Eq. (S1).

The pharmacological meanings of S-Eq constants were described as follows. ①Typical S-curve exhibited a symmetric figure centered on 50% y_max_ (y_50_). The dose corresponding to y_50_ is ED_50_ in pharmacology. S-curve was divided by y_50_ into low and high two parts. Each part had an inflection point located about 90% y_max_ (y_90_) and 10% y_max_ (y_10_), *i.e*., ED_90_ and ED_10_. There was a near straight line between the two inflexion points. The y_50_ site was depended on constant b and k. The morphological features reflect drug reaction law. ②The y_max_ and a value determined S-curve height. y_max_ = 1/a. S-curve height was high when the y_max_value was large. S-curve height was low when a value was large. However, the size of y_max_ and a value unaffected S-curve shape, such as curves [2 and 4]. ③k, affinity constant or S-curve slope, depends on the ratio of products to reactants (Eqs. A3 and A7-A9). Smaller doses produced larger effects at larger k values, such as curve [9]. Conversely, larger doses produced smaller effects at smaller k values, such as lines [3, 4, 6, and 7]. A straight line occurred in a wide range when k<0.1, such as line [7]. ④ Constant b determine intercept size. Intercept (y_0_) was the effect value at *x*=0. y_0_=1/(a+b). When a and k were determined, the bigger the b value was, the smaller the y_0_ value was, such as curves [1-4 and 9]. Conversely, the smaller the b value was, the bigger the y_0_ value was, such as curves [5-8 and 10]. S-curve shifted to the left with the b value decrease, such as curve [10].

In enzyme-substrate reaction, no effect occurs in the absence of enzymes when substrate is stable, *i.e*., y_0_=0 at *x*=0. At *x*=0, *y*≠0 showed that substrate was unstable or self-decomposition, y_0_ is the exhibition of substrate performance.

The effect indicators produced by drug reaction is usually expressed with DR, substrate surplus ratio (SR), or GR^18-21^. C_T_, C_D_ and C_S_ represent the concentrations of total, depleted, and surplus substrate, respectively. Effects are expressed with y_d_ and y_s_. The relationship among them is as follows.

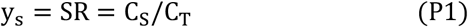

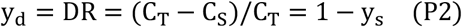

C_T_ are setting value. C_S_ and C_D_ are detection values. y_s_ and y_d_ are inverse functions of each other. The meaning of y_d_ and y_s_ are the same as the y meaning in Eq. (S1). The pharmacological meaning of constants (a, b, and k) can be reflected by effect *y*.

Based on y_max_ =1/a, the size of a determined by y_max_, clearly, y_max_ is not detection value because *x* value is impossibly set as ∝. When *x* =0, y_d,0_ has minima, and y_s,0_ has maxima. y_d,0_ = y_s,max_. y_d_ and y_s_ are both functions of C_T_ and C_S_. C_T_ is the setting value. C_S,0_ is the blank test values related to substrate properties. The y_max_ and a value can be determined by the y_d,0_ calculated with C_T_ and C_S,0_ (see the example below).

The meaning of constant b is expressed by intercept y_0_. y_0_ =1/(a+b). y_0_ is equivalent to the blank experiment value, reflecting substrate performance. The b value can be determined following y_0_ and a being solved.

Constant k, affinity constant, is a key parameter reflecting DRP. The k value can be calculated by putting a value and b value into Eq. (S1), but it is usually solved by S-Eq regression.

### 2.3. The DRP shown by S-curve shape

The purpose of pharmacological experiment is to test DRP. Constants (a, b, and k) were unable intuitively to express DRP. S-curve was divided by the two inflexion points into anterior, middle, and posterior parts as marked by the effect value of y_10_, y_50_, and y_90_. Their corresponding dose parameters (ED_10_, ED_50_, and ED_90_) directly express DRP (Table 1). Anterior part showed the transition of reaction status in test system from low efficiency to high efficiency. Driven by MAL, the effect in middle part proportionally increased with dose rise. Posterior part showed that drug reaction speed began slowing down, and effect no more proportionally increased with dose rise, which is related to excessive drugs, products, and the interaction of reactants and products. The effect value below 90% y_max_ at posterior inflexion point (y_90_) indicated that dose density reached saturated dose (SD). Notably, the effect value at saturated doses was not on S-curve and should not be included in the analysis data of PERs.

**Table 1.** Relationship between ED_50_ and constants.

| No* | Constants | ED<sub>10</sub> | ED<sub>50</sub> | ED<sub>90</sub> |
| --- | --- | --- | --- | --- |
| [1] | a=1, b=10, k=1 | 0.1054 | 2.3026 | 4.4998 |
| [2] | a=2, b=10, k=1 | -0.5878 | 1.6094 | 3.8067 |
| [3] | a=1, b=10, k=0.5 | 0.2107 | 4.6052 | 8.9996 |
| [4] | a=2, b=10, k=0.5 | -1.1756 | 3.2189 | 7.6133 |
| [5] | a=1, b=1, k=1 | -2.1972 | 0.0 | 2.1972 |
| [6] | a=1, b=1, k=0.5 | -4.3945 | 0.0 | 4.3945 |
| [7] | a=1, b=1, k=0.1 | -21.972 | 0.0 | 21.972 |
| [8] | a=1, b=3, k=1 | -1.0986 | 1.0986 | 3.2958 |
| [9] | a=1, b=10, k=4 | 0.0263 | 0.5756 | 1.1250 |
| [10] | a=1, b=0.2, k=1 | -3.8067 | -1.6094 | 0.5878 |

\* same as Fig. 2. ED<sub>10</sub>, ED<sub>50</sub> and ED<sub>90</sub> were calculated with constant values entering Eq. (S1).

ED_10_, ED_50_ and ED_90_ in Table 1 were calculated with constants values (Fig. 2). ED_i0_ size depends on constant values. Low ED_50_ shows that drugs at low doses can produce high effect, indicating that drug performance is good. Otherwise, high ED5_0_ indicates that the drug performance is poor.

## 3. The S-curve analysis method

### 3.1. The solving method of S-curve

The purpose of pharmacological experiments is to understand DRP by analyzing dose-effect relationship. The analysis method of PERs is generally to plot dose (*x*_*i*_) and effect (*y*_*i*_) curves. However, how to analyze dose-effect curve (DEC), everyone has different view and methods^11,12,18^. The present study suggests that PERs should be analyzed as S-curve or its part in order to get more true drug reaction law.

An essential way to solve S-curve is to take *x*_*i*_ and *y*_*i*_ values into S-Eq to form a ternary first order Eq group. After the value of constant (a, b, k) is solved, drug performance parameters are calculated by substituting constant values into original Eq. The method solving the ternary first order Eq group has been not found in pharmacological study. Generally, according to DEC exhibited by PERs, appropriate mathematical formula is employed, and fitted into linear Eq. Drug performance parameters are calculated after the constant is determined by regression method. Under different experiment conditions, anterior, middle, and posterior part of S-curve can be exhibited by PERs. Anterior part was similar to exp curve, middle part similar to straight line, and posterior part similar to lg curve. Anterior part plus middle part or middle part plus posterior part was similar to hyperbolic curve. Accordingly, the partial exhibition form of S-curve, such as exp, lg, or Michaelis-Menten (M-) Eq, has been widely used in analysis of PERs. These methods belong to the approximate analysis methods of S-curve. For example, exp Eq directly ignores constant a in S-Eq. Eq. (S1) is transformed into exp Eq. (S3).

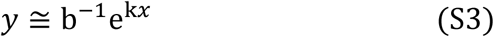

When r=1/a is put into Eq. (S2), exp Eq. (S4) forms by approximate transformation.

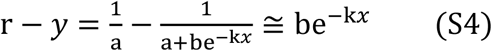

Eqs. (S3 and S4) are transformated by natural logarithm (ln) into linear Eqs. (S5 and S6).

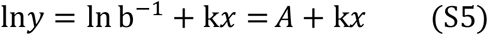

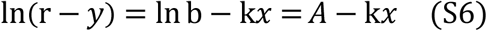

### 3.2. The S-curve linear fitting method

Analyzing S-curve requires to converting S-Eq into linear Eq. The linear fitting method was as follows. Eqs. (S1 and S2) are first converted into reciprocal function Eqs. (S1i and S2i), then transformed into exp Eqs. (S7 and S8) following the shift of constant a from equation right to left. Eq (S8) is reverse function of Eq. (S7).

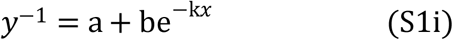

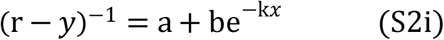

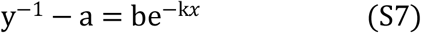

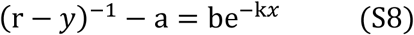

Take r=1/a into Eq. (S8) left to get the Eq. (*r* − *y*)^−1^ − a = a^2^/(*y*^−1^ − a).

Replace its denominator (*y*^−1^ − a) with the right of Eq. (S7), be^−k*x*^, to get Eq. (S9).

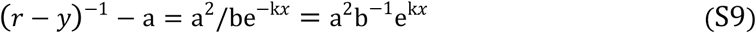

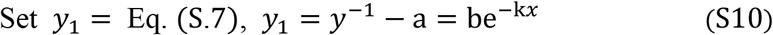

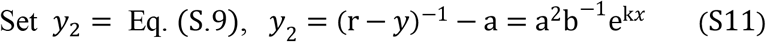

Following natural logarithm (ln) transformation of Eqs. (S10 and S11), linear fitting Eqs. (S12 and S13) are established. Eqs. (S12 and S13) are essential Eqs to solve S-curve.

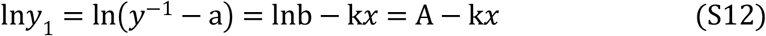

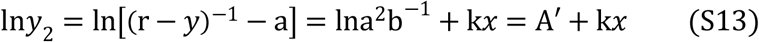

### 3.3. The linear fitting degree test of S-Eq

Conventional method of the test is to calculate the correlation index (R^2^) value by Eq^22^.

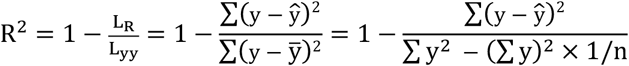

Here, y, ŷ, 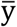, and n are effect value for regression calculation, expected true value of effect y, the mean value of effect y, and sample numbers, respectively. Below is to take S-curve y_1_ (a=1, b=9, k=1) in Fig. 1 as an example. Following *x*_*i*_ values entering original Eqs. (S1 and S2), the corresponding effect values (*y*_*i*_) are calculated and listed in lines 1 and 4 in (Table 2), respectively. Reqs. (S.14-S.17) are got when *x*_*i*_ and *y*_*i*_ are taken into the corresponding linear Eqs. (S5, S6, S12, and S13), respectively. The *y*_r*i*_ value reversely calculated by Reqs. (S14-S17) is listed into lines 2, 3, 5, and 6 in Table 2, respectively.

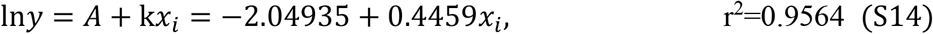

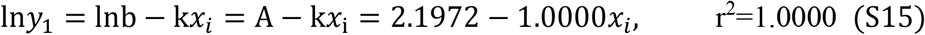

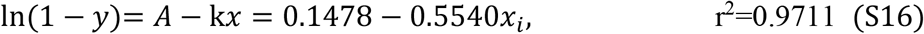

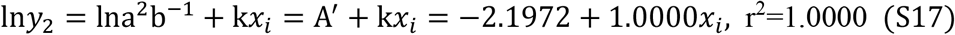

**Table 2.**
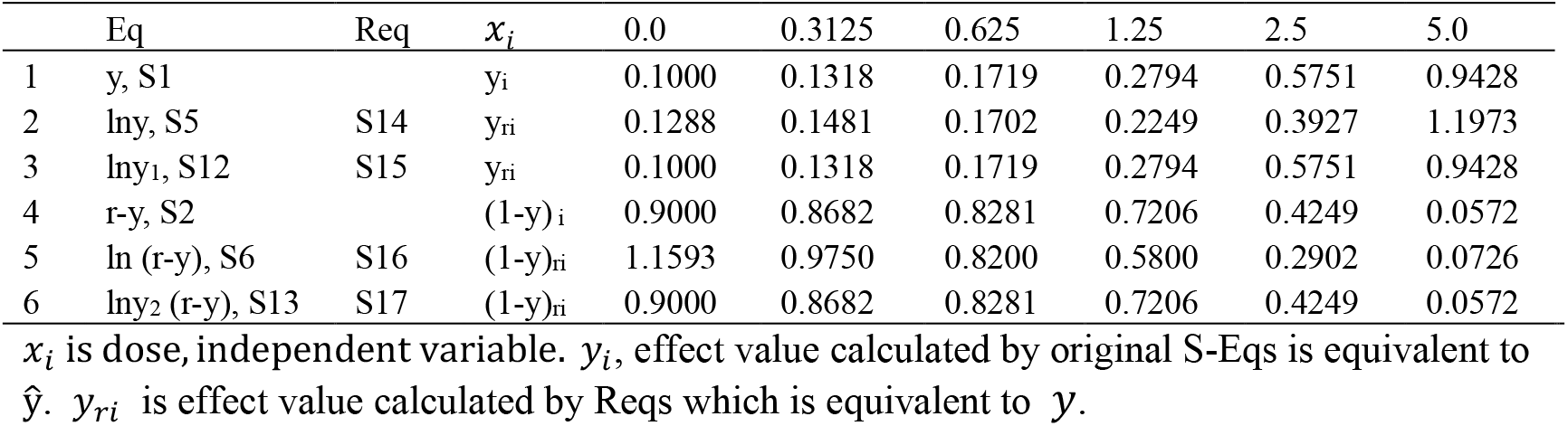
Analysis of R2 for the goodness-of-fit test.

L_R_, L_yy_ and R^2^ values are respectively calculated below, when *yi* values from S-Eqs. (S1 and S2) serve as ŷ and *y*_*ri*_ values from Reqs. (S14-S17) as *y* values. The results show that the linear fitness of S-Eq was 100%, whereas the linear fitness of exp Eq was only about 81%.

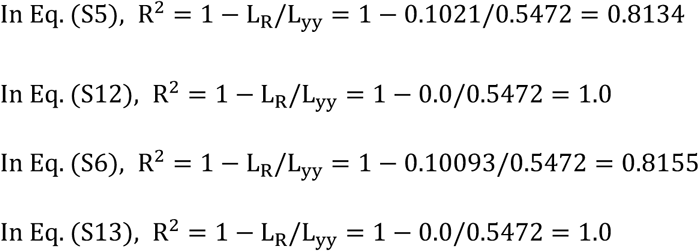

## 4. The example for application of the S-curve analysis method

The S-curve analysis method of PERs is based on linear fitting Eqs. (S12 and S13). The experimental results expressed by doses (*x*_*i*_) and effect values (*y*_*i*_) are taken into Eq. (S12 or S13) to carry out regression calculation and to obtain a, b, and k values. Then, the constant values are taken into Eq. (S1, S2, S12, or S13) to solve experimental parameters, such as ED_50_ and so on. Following it is to demonstrate the application of the S-curve analysis method by borrowing a set of data in our cytotoxic experiment (lines 1 and 2 on the left of Table 3).

**Table 3.**
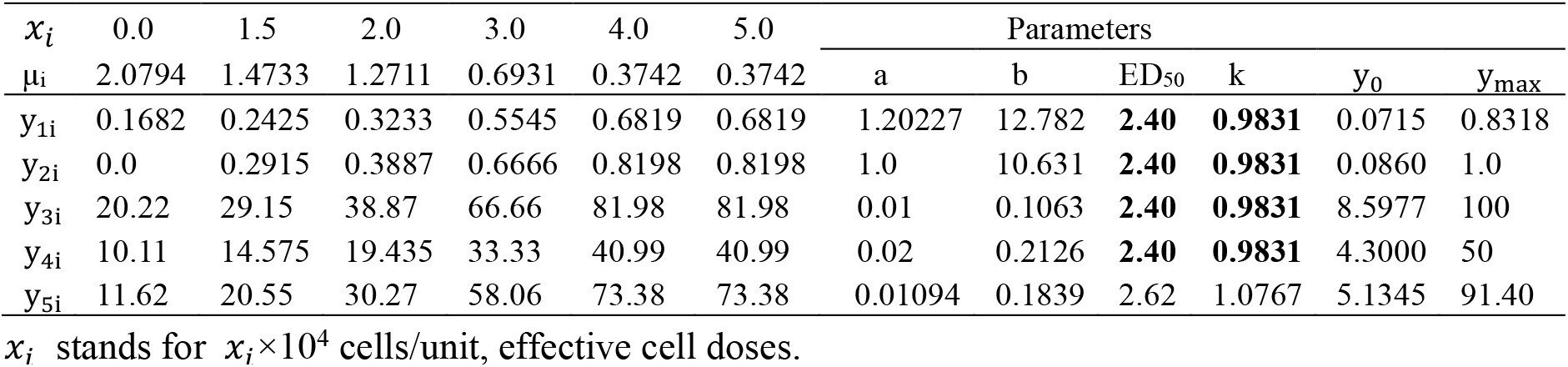
Cytotoxic experimental data and the parameters calculated by Reqs.

In the experiment, peripheral blood mononuclear cells (PBMCs) were used to effective cells equated to toxic drugs. Human chronic myeloid leukemia cell line (K562 cells) was used as target cells equated to substrate. Killing ratio (KR) of the target cells killed by effective cells was used as a cytotoxic effect indicator. KR = y_d_. y_d_ is a function of effect cell doses (*x*_*i*_). Target cell changes are detected by the limiting dilution cell clone forming inhibition assay and converted into KR by Eq. (P2-1). One living target cell forms one clone^23^. There were about 50 clone units per dose (*x*). The clone mean per dose unit is the mean (µ_i_) of viable target cells. The reactants in the experiment consist of effective cells and target cells, and the products are the reduction value of target cells. Although effective cells do not necessarily die after the interaction with target cells, they cannot form products. So, there is only one product in the experiment. According to Eq. (9), the effect *y* = KR. KR per dose were respectively calculated when configured target cell density (μ_20_=2.5 cells/unit) or tested target cell clone mean at *x*=0 (μ_20_=2.0794 cells/unit) was used as *substrate* (effect conversion factor). The two µ_**0**_ were employed mainly because both cases exist. Actually, only one of both is used.

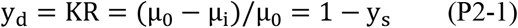

Set y_1i_=KR_1i_ and y_2i_=KR_2i_ (Table 3). According to y_d.0_ = y_s.max_,

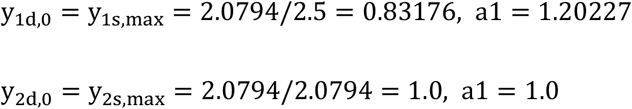

Reqs. (S18 and S19) are got when a_1_ and a_2_ values are put into Eq. (S12) to do regression, respectively. Notice, in this example, the effect value of y_d,0_ and y_5.0_ corresponding to *x*=0 and *x*=5.0 were not listed into the regression. The reasons are as follows. (1) The y_d,0_ value at *x*=5.0 is the blank experimental value. There are two cases for using µ_10_ as *substrate* if y_d,0_ = 0.1682 is put into regression. (1.1) KR is detected by the limiting dilution cell clone forming inhibition assay. There are only target cells in the blank experimental unit. There are target cells and effective cells in the experimental unit. The non-cytotoxic cells in the effective cells, as nutritive cells, can assist in target cell clone formation, so that the clone formation ratio in the clone unit with nutritive cells can significantly be higher than that without nutritive cells. In the assay, the y_d,0_ value detected in the blank experimental unit was lower than the actual value. Thus, it was not included in the regression calculation. (1.2) The y_d,0_ value should be included in the regression calculation if the blank experimental group contains the proper number of nutritive cells, such as red blood cells. However, this example is the previous case. (2) If µ_20_ is used as *substrate* (Eq. P2-1), then y_d,0_ = 0. y_d,0_ = 0 entering Eq. (S12) is meaningless, which is caused by the rate conversion with µ_20_. (3) The y_5.0_ value (0.6819) at *x*=5.0 is a saturation density value in this example. ED_90_ calculated by Req. (S18) is at the position of *x*=4.6394. Then, y_90_=0.7486 and y_5.0_=0.7716 at *x*=5.0. Computationally, the y_5.0_ value (0.7716) is greater than the y_90_ value (0.7486). However, the y_5.0_ value tested (0.6819) in Table 3 is smaller than the y_90_ value (0.7486). So, the y_5.0_ value tested belongs to the saturation density value, and is not included in the regression calculation (Section 2.3).

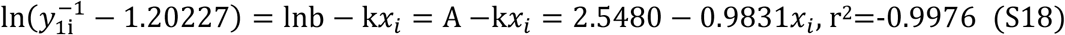

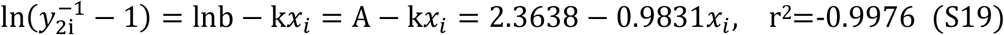

The y_max_ value is from 0.83176 to 100, and a_3_=0.01 when *y*_3i_ is 120.227 times of *y*_1i_. The y_max_ value is from 0.83176 to 50, and a_4_=0.02 when *y*_4i_ is 60.1135 times of *y*_1i_. Reqs (S20 and S21) are got when a_3_ or a_4_ is put into Eq. (S12) to do regression, respectively.

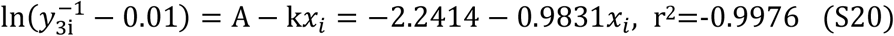

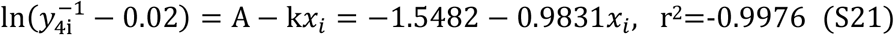

The following DRP are calculated by Reqs. ED_50_ can be respectively obtained when y_50_ (y_max_×0.5) is put into Reqs. (S18-S22). The y_0_ value can be respectively obtained when *x*=0 is put into Eqs. (S18-S22). The constant b value can be got by lnb = A in Eq. (S.12). The data is shown on the right in Table 3.

The substrate effect on PERs is mainly expressed by the y_d,0_ value, and also expressed by y_0_ value (S-curve intercept). y_d,0_ is the experimental value at single dose in the absence of effective cells, whereas y_0_ is the experimental value calculated by regression at multiple-point doses in the presence of effective cells. Clearly, y_0_ can better reflect the substrate effect on PERs than y_d,0_. Thus, y_d,0_ is not used by the S-curve analysis method in regression calculation. The substrate effect is eliminated by the S-curve zeroing method that S-curve wholly decrease by one y_0_. That is to say, the y_i_ - y_0_ value is taken into Eq. (S12) to do regression calculation when y_0_ is solved by Reqs. For example, Eq. (S22) is the y_5i_ Req got after y_3i_ curve zeroing. y_3,0_=8.5977. The results are listed in the bottom row of Table 3.

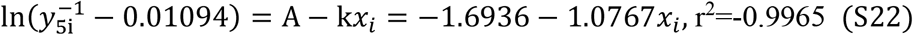

The above analysis can be summarized as follows (Table 3 and Fig. 3A). ①The y_max_ and a value can be got by Eq. (P2-1) when µ_10_ (configured density) or µ_20_ (the blank experimental value) is used as *substrate* to perform the rate conversion. The experimental parameters are calculated after Reqs. (S18-S21) are obtained using the data in Table 3. Different *substrate* can produce the different value of y_max_, a, b, and y_0_. However, the k and ED_50_ value remains unchanged as long as the experimental value is unchanged. The same k and ED_50_ value from y_1i_ to y_4i_ in Table 3 show that the different *substrate* has no effect on the analysis result, revealing that k and ED_50_ are the key parameters to reflect DRP. ②Magnification or deflation of the y_max_ and a value do not affect analysis results. ③The S-curve zeroing method should more truly reflect DRP than the method of subtracting the y_d,0_ value as far as the substrate effect on PERs is concerned.

**Figure 3.**
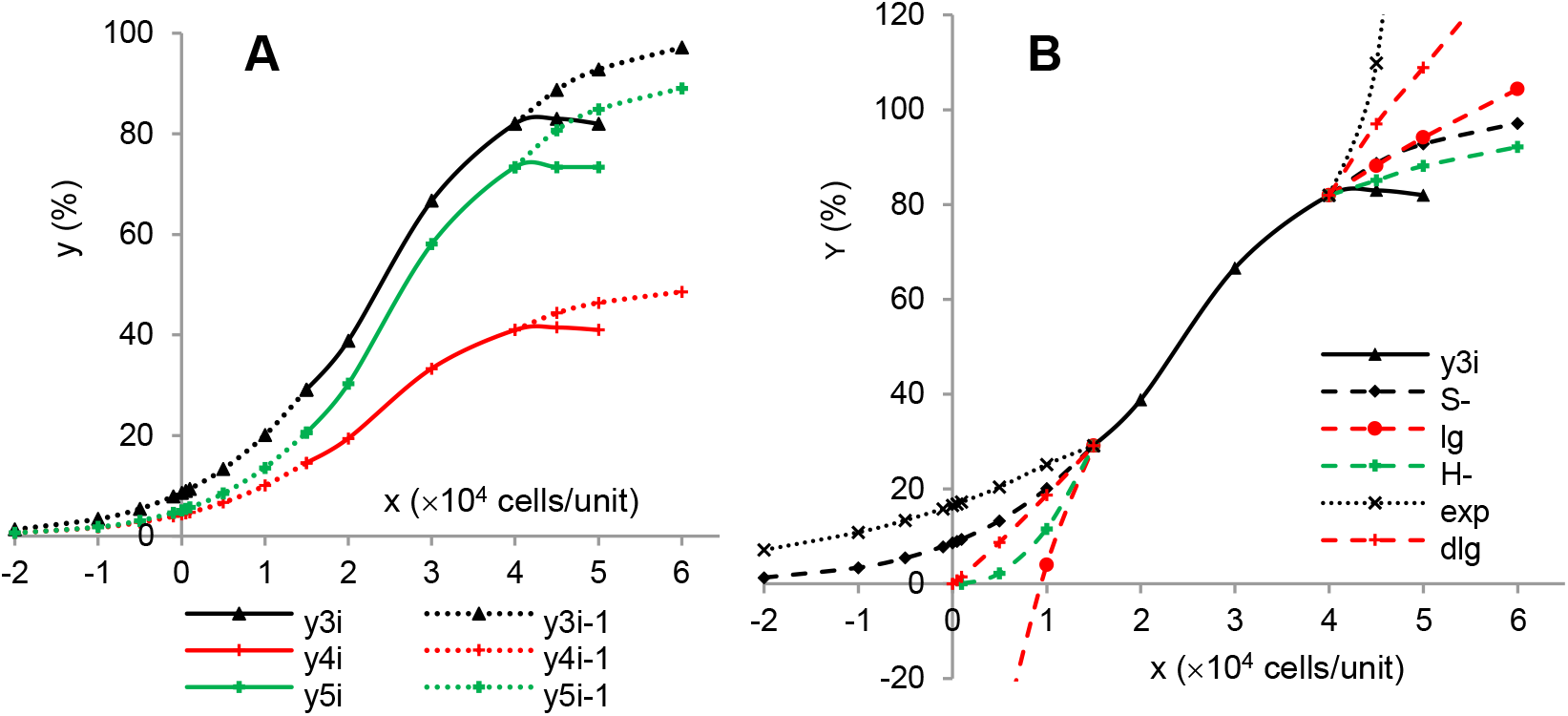
The DEC in the cytotoxic experiment (A) and the comparison of analysis results on y_3i_ (B). Solid line is experimental values. Dashed line is curve extension parts calculated by the Reqs in order to show S-curve shape. y_4_ curve is 50% lower than y_3_ curve in position. The y_5i_ curve is y_3i_ curve zeroing.

## 5. The comparison of five analysis methods in PERs

Up to now, traditional PERs have been usually processed with the corresponding Eqs, such as exp, lg, dlg and hyperbolic Eq, according to the curve shape exhibited by PERs. Experimental parameters are solved by regression following linear fitting of the corresponding Eqs. The follow was to analyze and compare y_3i_ results in Table 3 by the below five methods. ①The S-curve analysis method used Eq. (S12). ②The exp curve method used approximate fitting Eq. (S5). ③The lg curve used *y* = A + kln*x* (lg-1) ④The dlg curve Eq used ln*y* = A + kln*x* (lg-2). ⑤Hyperbolic Eq derived from Hill Eq. (H-Eq) and M-Eq.

### 5.1. Basic H-Eq form

H-Eqs. (H1-H2) are formed based on receptor-ligand reaction^4,24^. Eq. (H3) is got when k_d_ is put into Eq. (H1)^24^.

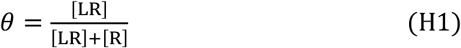

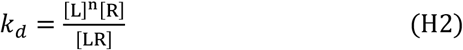

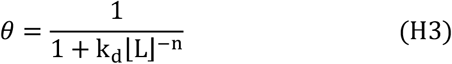

Here, k_d_, n, Θ, [L], [R], and [LR] are apparent dissociation constant, Hill coefficient, effect parameter, ligand, receptor, and ligand-receptor conjugate concentration, respectively. Replace Θ with *y*, [L] with *x*, and k_d_ with b, *i.e*., Θ=*y*, [L]=*x*, and b=k_d_, then Eq. (H3) is converted into Eq. (H4).

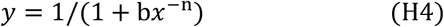

Eq. (H4) can be converted into Eqs. (H5 and H6) by inversion, shifting of 1 to left, and taking logarithm.

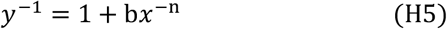

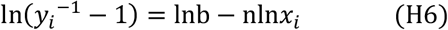

Here, Eq. (H5) shows that H-Eq is power function. H-curve shape is determined by power n (H coefficient) and constant b. H-curves with different n and b values are all hyperbolic curve (Fig. S1), indicating that Hill curve is really not sigmoid curve. In fact, H-Eq has been employed in sigmoid curve^7,25,^.

[LR]/[R] is equivalent to GR. According to the definition of Θ in Eq. (H1), Θ is not equivalent to GR, such as (H7). To put *Θ* = *y* = 1/[1 + (*GR*)^−1^] into Eq. (H5), Eqs. (H8-H10) are got. Eq. (H10) is equivalent to Eq. (H6), but its effect meaning is different.

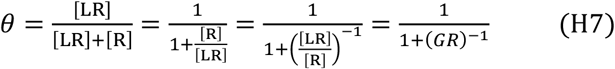

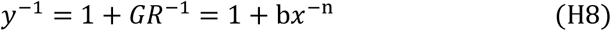

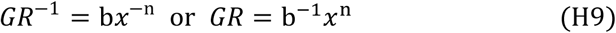

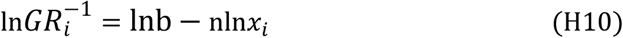

### 5.2. Basic M-Eq form

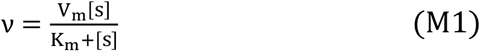

Here, V_m_ is maximal velocity as an effect indicator^26,27^. The core of M-Eq is to define M constant (K_m_) as the dose value when ν = V_m_/2. That is, K_m_ is equivalent to ED_50_. Eq. (M1) is written into Eq. (M2).

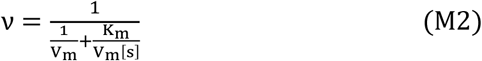

Eq. (M2) is converted into linear Eq. (M3). Eq. (M4) is got when Eq. (M3) is multiplied by V_m_.

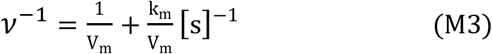

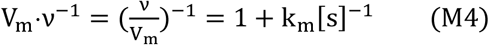

M-Eq expresses the relationship between ν and reactant dose [S]. Obviously, M-Eq is derived from MAL defined by van’t Hoff. ν/V_m_ is equivalent to GR. To replace ν/V_m_ with *y*, [S] with *x* and K_m_ with b (*y*=ν/V_m_, *x*=[S], and b=K_m_), Eq. (M4) is converted into Eqs. (M5-M7).

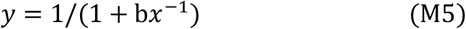

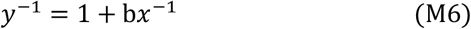

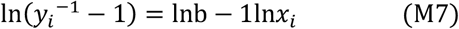

Like Eq. (H5), Eq. (M6) is also power function. Clearly, Eq. (M6) is a special case of Eq. (H5) at n=1. It is hyperbolic curve located in quadrants I and III with 0 point as the symmetric center (Fig. S1). Eqs. (H6 and M7) are both dlg function Eqs.

### 5.3. The experimental results processed by the below methods

The following uses the above method to obtain Reqs. The data is from y_3i_ in Table 3. Experimental parameters are listed in Table 4.

**Table 4.** The parameters calculated by five methods

| Curve | $R^2$ | $ED_{50}$ | $k$ | $x$ domain | $y_0$ | $y_{max}$ |
| --- | --- | --- | --- | --- | --- | --- |
| S- | 0.9976 | 2.40 | 0.9831 | $-\infty < x < \infty$ | 8.60 | 100% |
| Exp | 0.9789 | 2.63 | 0.4209 | $-\infty < x < \infty$ | 16.52 | 100% |
| lg | 0.9936 | 2.27 | 56.00 | $x \neq 0$ | $x \rightarrow -\infty$ | 100% |
| dlg | 0.9949 | 2.46 | 1.0945 | $x \neq 0$ | $x \rightarrow 0$ | 100% |
| H11 | 0.9948 | 2.46 | 1.0945 | $x \neq 0$ | $x \rightarrow 0$ | 100% |
| H12 | 0.9934 | 2.25 | 2.5049 | $x \neq 0$ | $x \rightarrow 0$ | 100% |
| M8 | 0.9934 | 2.25 | 2.5049 | $x \neq 0$ | $x \rightarrow 0$ | 100% |

$y_0$ , control effect. $y_{max}$ , maximal effect.

#### 5.3.1. *Reqs derived from the S-, exp, l*g, and *dlg curve method*

① S-Eq from Eq. (S12): 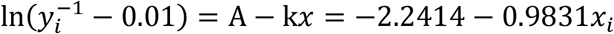, r^2^=-0.9976
② exp Eq from Eq. (S5): ln*y*_*i*_ = *A* + k*x* = 2.8048 + 0.4209*x*_*i*_, r^2^=-0.9789
③ lg Eq from Eq. (lg-1): *y* = A + kln*x* = 3.9917 + 56.0045ln*x*_*i*_, r^2^=-0.9936
④ dlg Eq from Eq. (lg-2): ln*y*_*i*_ = A + kln*x* = 2.9291−1.0945ln*x*_*i*_, r^2^=-0.9949

#### 5.3.2. Regs derived from the hyperbolic curve method

In the calculation method of hyperbolic curve, n in Eq. (H6) and 1 in Eq. (M7) as set symbol are just different in form. Their size can be n, 1 or number else depended on experimental results^24, 28^. When experimental data is put into Eqs. (H6 and M7) their Reqs are below got to calculate performance parameters, such as ED_50_, K, K_d_, K_m_, and *y*_0_. Approximate number (0.0001) is substituted for 0 when the *y*_0_ value is calculated because it is impossible to let *x*=0 in H-Eq. Req. (H11) is got in H-Eq with *Θ* = 1/[1 + (*GR*)^−1^]. The same Reqs. (H12 and M8) are got in H-Eq and M-Eq with original definition of Θ and M, respectively. Their parameters were calculated (Table 4).

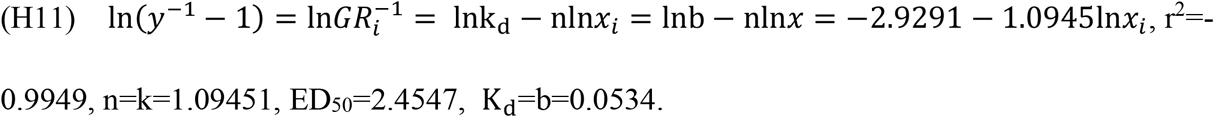

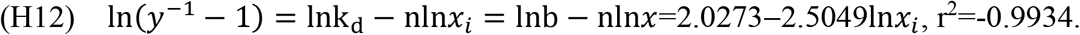

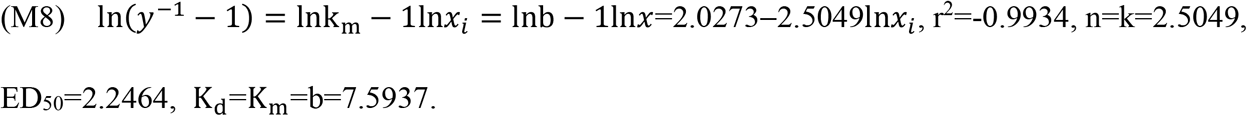

Table 4 and Fig. 3B show that the curve exhibited by the same set of experimental data is actually a part of S-curve. It can also be considered as a part of exp, lg, dlg, and hyperbolic curve. Based on different understanding of the curve, different results were got when different analysis methods were employed. The meaning of experimental parameters (ED_50_, k, *y*_0_, y_max_) expressed by the S-curve analysis method is very clear. The present method can more truthfully reflect drug performance. The result obtained by the exp curve method was close to S-curve in low dose range and deviated greatly in high dose range. Unlike the exp analysis, the result got by the lg curve method were close to S-curve in high dose range and deviated greatly in low dose range. The results got by the dlg curve method were approximately a straight line. The results got by the hyperbolic curve method and the S-curve method deviated far from each other. This reason is as follows. Theoretically, Θ as the effect form set by H-Eq is inconsistent with the form defined by MAL. To replace Θ with GR, the Reqs obtained from the experimental results are the same as those obtained with the dlg curve. M-Eq is derived from MAL defined by van’t Hoff. As mentioned in Section 2.1, it is unreasonable to use ν as the effect indicator. The meaning of ν/V_m_ and [LR]/[R] is also different. Importantly, the dose-effect relationship shown by PERs is actually not hyperbola but S-curve.

The above result shows that the dose effect relationship exhibited by PERs is regarded as S-curve to be analyzed, which is both reasonable and can truly reflect drug reaction law. In the traditional analysis methods, the part pattern exhibited by PERs is regarded as a whole pattern and analyzed by the approximate mathematical model, which cannot accurately reflect drug reaction law.

## 6. Conclusion

The S-curve analysis method for PERs is based on linear fitting method of S-Eq with 100% fitness. Additionally, S-curve shape and S-Eq constants (a, b, and k) are definite in pharmacological meaning. The experimental parameter obtained from the present method can be closer to real DRP than those from the traditional methods. Therefore, the elucidation of the S-curve analysis method has important significance for pharmacological and biomedical study.

## Supporting information

Supplemental figgure 1

## Abbreviations

b: a major factor of intercept
C_T_: total concentration
C_D_: depleted concentration
C_S_: surplus concentration
DEC: dose-effect curve
ED_50_: the dose corresponding to 50% y_max_
DR: depletion ratio
DRP: drug reaction performance
dlg: double logarithmic
exp: exponential
H-Eq: Hill equation
GR: growth ratio
*k*: reaction velocity constant, affinity coefficient
Kd: apparent dissociation constant
KR: Killing ratio
K_m_: Michaelis-Menten constant
lg: logarithmic
ln: natural logarithm
MAL: Mass action law
M-Eq: Michaelis-Menten equation
PER: pharmacological experiment result
Req: regression equation
SD: saturated dose
S-curve: sigmoid curve
S-Eq: sigmoid curve equation
SR: surplus rate
V_m_: maximal reaction velocity
*x*: dose, independent variable
*y* (effect): a function of *x* or dependent variable
y_max_: maximal effect
y_0_: S-curve intercept.

## Acknowledgments

This work was supported by the National Natural Science Foundation of China (No. 81573820).

## Supplementary information

Supplementary information can be found in Figure S1.

## Author contributions

Chengyan Zhao: Conceptualization, investigation, writing - original draft. Qingxia Niu: investigation, Writing - review & editing.

