## Supplemental figgure 1 for "A sigmoid curve analysis method for pharmacological experimental results"


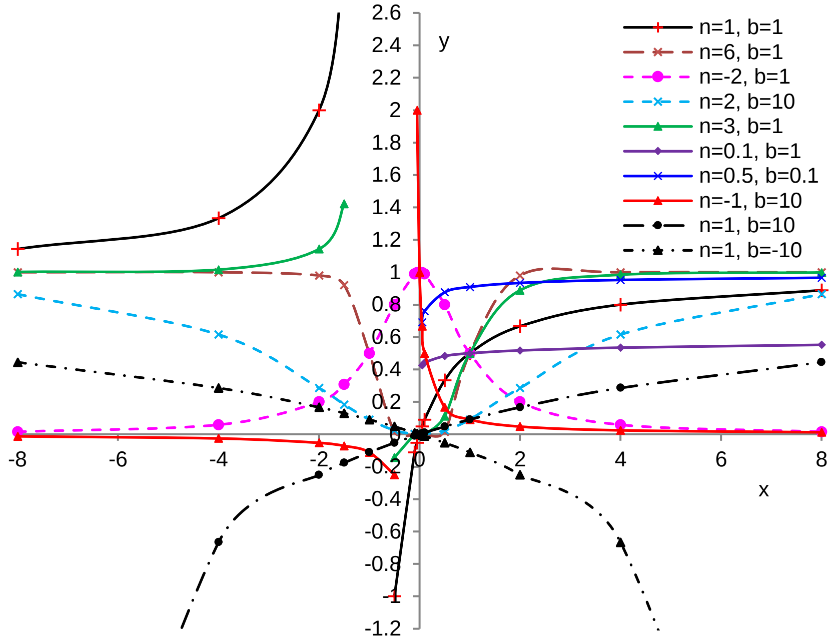


**Figure S1** Hill curves based on H-Eq. (H4). Here, n and b stand for power and constant, respectively. Different H-curves are all hyperbola with the change of n and b values. H-curve at n>0 is centrally symmetric when n=odd number, and is axis-symmetric when n=even number. The large the n value is, the large the curvature of H-curve is. Generally, there is no curve in quadrant 4 for power function unless b<0.
